# Quantitative Genetic Scoring, or how to put a number on an arbitrary genetic region

**DOI:** 10.1101/2020.12.15.422886

**Authors:** GH Schoenmacker, P Vlaming, J Pallesen, MY Pikulina, AH Ghamarian, D Demontis, A Børglum, TE Galesloot, G Poelmans, B Franke, T Claassen, T Heskes, JK Buitelaar, A Arias Vásquez

## Abstract

**Motivation:** With the increasing availability of genome-wide genetic data, methods to combine genetic variables with other sources of data in statistical models are required. This paper introduces quantitative genetic scoring (QGS), a dimensionality reduction method to create quantitative genetic variables representing arbitrary genetic regions.

**Methods:** QGS is defined as the sum of absolute differences in the genetic sequence between a subject and a reference population. QGS properties such as distribution and sensitivity to region size were examined, and QGS was tested in six different existing genomic data sets of various sizes and various phenotypes.

**Results:** QGS can reduce genetic information by >98% yet explain phenotypic variance at low, medium, and high level of granularity. Associations based on QGS are independent of both size and linkage disequilibrium structure of the underlying region. In combination with stability selection, QGS finds significant results where a traditional genome-wide association approaches struggle. In conclusion, QGS preserves phenotypically significant genetic variance while reducing dimensionality, allowing researchers to include quantitative genetic information in any type of statistical analysis.

**Availability:** https://github.com/machine2learn/QGS

**Supplemental information:** Supplemental data are available online.

## 1 Introduction

The rise of affordable, high throughput genotyping and next generation sequencing techniques in the past decade has made whole genome genotyping and sequencing affordable options for biomedical research. With increasing sample sizes and thus statistical power, genome-wide association studies (GWAS) have effectively replaced candidate gene studies for complex multifactorial conditions (Duncan *et al.*, 2019). These GWAS show through estimation of variant-wide explained variance that genetics can explain a significant portion of variance in these complex, multifactorial phenotypes, ranging from easily measurable traits such as height and body mass index (BMI) (Yengo *et al.*, 2018) to complex psychiatric disorders such as schizophrenia (Pardiñas *et al.*, 2018) and attention deficit/hyperactivity disorder (ADHD) (Grigoroiu-Serbanescu *et al.*, 2020).

The growing availability of genome-wide genetic information combined with its proven ability to explain variance in complex phenotypes further fuels the wish to combine genetic variables with other outcomes (e.g. neuroimaging or behaviour) in building disease-explanatory statistical models. However, methods to transform measured genetic values into meaningful numeric variables are currently lacking.

This paper introduces a new method to summarise genetic information: quantitative genetic scoring (QGS). QGS is a genetic dimensionality reduction method to create quantitative genetic variables representing arbitrary genetic regions. In contrast to the GWAS based summary statistics, QGS is phenotype-agnostic and therefore does not depend on previous association results. Our approach estimates a numeric value corresponding to the observed allelic variation of arbitrary genetic regions selected by the user (exons, genes, gene sets, genetic pathways, intergenic regions) irrespective of allele frequency and impervious to the size and linkage disequilibrium patterns of the selected region. The estimated QGS can be used in any downstream statistical analysis approach as a quantitative variable. Because it can drastically reduce the number of variables in genome-wide analyses while retaining genetic information, it also allows for the use of more computationally complex genome-wide methods.

The goal of this paper is to introduce QGS as a method to summarise genetic variation within a region into a numeric value. We show that QGS retains genetic information while reducing dimensionality. Software to calculate QGS scores working with PLINK-format (Purcell *et al.*, 2007) or Variant Call Format (VCF; Danecek et al., 2011) data is available at https://github.com/machine2learn/QGS.

## 2 Materials and Methods

### 2.1 QGS

While not categorically true, in genetics the rarer allele (i.e. the minor allele) often is associated with increased disease risk instead of protection. This holds for rare mutations (Kryukov *et al.*, 2007) as well as common variants (Park *et al.*, 2011). Continuing this reasoning, QGS extends the concept of population similarity beyond single variant MAF to whole genetic regions.

QGS assigns a numeric value 0 ≤ QGS ≤ 1 to any arbitrary genetic region. The QGS is based on the average difference between an individual’s genetic information (in the form of genotypes) and that of a reference population. Intuitively, QGS can be interpreted as a measure of genetic “distance” compared to the reference population: a low score indicates higher similarity to the reference population, whereas a high score indicates a lower similarity.

To compute the QGS the following structures are defined:

- An arbitrary genetic region defined by N_var_ number of variants;
- A sample individual genetic dosage (or hard calls) information vector s ={s_1_,…, s_Nvar_} where each element represents genetic dosage data (or hard calls) 0 ≤ s_i_ ≤ 2, i.e. the (probabilistic approximation of the) minor allele count for variant ¿;
- A reference population matrix of N_refs_ individuals containing genetic dosage (or hard call) data:

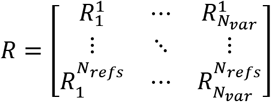

Finally, the QGS itself is defined as the grand sum of the absolute difference between *R* and s, scaled to 0-1 by dividing by the total number of alleles in the reference population:

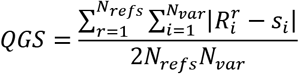

QGS was implemented in C++ and is available as a command-line program to compute QGS from VCF or PLINK files. An example of a QGS calculation is given in the supplement.

### 2.2 Related works

Below is a non-exhaustive overview of methods for collapsing genetic information into a numeric construct. For common variants, methods exist for combining summary statistics from single nucleotide polymorphisms (SNPs), i.e. effects sizes or p-values) into a single statistic using metaanalysis and/or permutation techniques (e.g. Huang et al., 2016; Liu et al., 2010; S. Purcell et al., 2007; Wang, Li, & Bucan, 2007). These methods provide a combined estimate for the effect or p-value, but do not create a numeric construct. One exception is MAGMA (de Leeuw *et al.*, 2015), which translates a genetic region into a vector of varying length using SNP-based principal components.

Another method, polygenic risk scoring (PRS; S. M. Purcell et al., 2009), does provide a single construct by multiplying the number of minor alleles with a previously found effect size for a certain phenotype in an independent population and summing the result across all variants. While PRS has proven ability to explain phenotypic variance in independent samples, like the meta-analysis techniques its use depends on the public availability of large-sample based GWAS results, because the explanatory power of PRS is proportional to the sample size of the original GWAS (Dudbridge, 2013).

Burden scores for rare variants attempt to quantify the genetic burden of such variants by dichotomising or counting of mutations (e.g. Asimit, Day-Williams, Morris, & Zeggini, 2012; B. Li & Leal, 2008; Morgenthaler & Thilly, 2007). Other rare variant association methods exist (Lee *et al.*, 2014), but burden scores are conceptually closest to QGS, because they also attempt to put an association-independent value to a genetic region. QGS differs from burden scores and unweighted PRS that count minor alleles by calculating a difference from a reference population and working with dosages. This allows QGS to work for both common as well as rare variants.

### 2.3 Data sets

In total, six data sets of different sizes and outcomes were used. A schematic overview of the genetic data sets and analyses can be found in Figure 1. The largest sample was the UK Biobank (UKBB) with the continuous psychiatric phenotype sociability (UKBB-s) (Bralten *et al.*,2019). The second largest sample was also from UKBB with the binary phenotype of lifetime cannabis use (UKBB-c) (Pasman *et al.*, 2018). Third was the Lundbeck Foundation Initiative for Integrative Psychiatric Research (iPSYCH) sample with the binary phenotype of ADHD status (Demontis *et al.*, 2019). Fourth was the Nijmegen Biomedical Study (NBS) with the continuous phenotype of body mass index (BMI) (Galesloot *et al.*, 2017). Fifth was the public 1000 Genomes Project phase three (1000G) (1000 Genomes Project Consortium *et al.*, 2015). Sixth was the public International HapMap Project phase three (HapMap3) (Frazer *et al.*, 2007). iPSYCH is a case/control cohort, whereas the others are general population cohorts. A more detailed description of the samples and data collection is given in the supplement.

**Figure 1:**
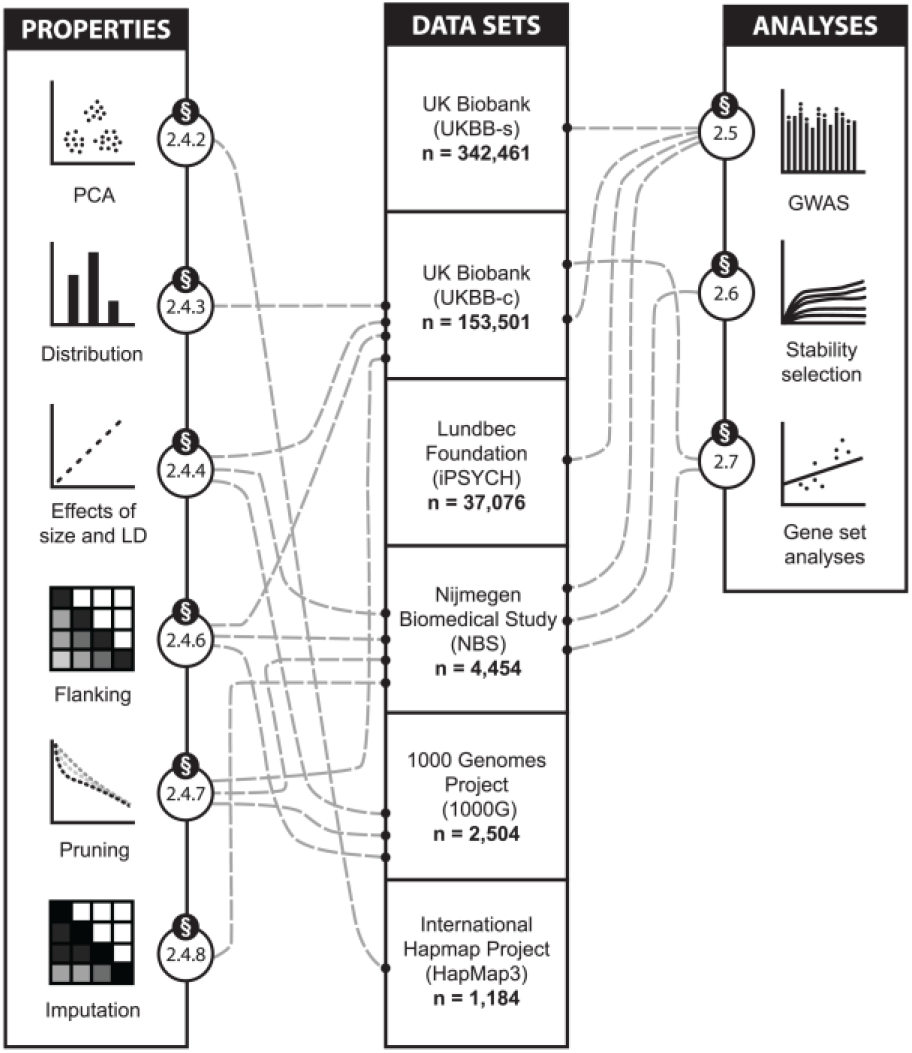
Schematic overview of the data sets and analyses. The six data sets used are shown in the middle in descending order of size. On the left-hand side, the QGS properties that were examined are displayed. On the right-hand side, the performed genetic analyses are shown. The section numbers in the circles (e.g. “§2.4.2” forPCA and “§2.6” for stability selection) refer to Methods sections. The lines link the data sets with the properties and analyses for which they have been used. PCA: principal component analysis; LD: linkage disequilibrium; GWAS: genomewide association study.

### 2.4 Analyses

The main goal of QGS is to quantify and summarise genetic information into a numeric value and make it available for downstream statistical analyses. To this end, it was important to verify that QGS indeed retains useful genetic information while reducing dimensionality. In addition, we have examined several properties of QGS, including its distribution, effects of region size and linkage disequilibrium structure (LD; correlation structure across genetic variants) on association, effects of including flanking regions of genes, pruning steps, and imputation quality control. Thirdly, QGS was compared to GWAS in different study designs (case/control and population-based) and for different phenotypes. Fourthly, stability selection, a computationally intensive method for variable selection, was applied with QGS. Lastly, the efficacy of QGS was shown in two gene set based associations.

#### 2.4.1 QGS Calculation

QGS values were calculated for all genes in all data sets using the European population of 1000G Phase 3 (n=503) as a reference population, except in the case of HAPMAP3 and UKBB which were used as their own reference. Gene regions were defined by the GENCODE gene annotation release 29 (Harrow *et al.*, 2012).

QGS can be calculated using, for example, only protein-coding genes but, because the gene-based QGS results only include gene regions, a large portion of the genome will not be covered. To gain coverage for intergenic regions, a block-based QGS was also calculated for NBS and UKBB with uniform blocks of 66kb length, which is the mean size of protein-coding genes in the above annotation. This was done to make block regions roughly comparable to gene regions in terms of size and information contents. A sliding window of 10kb was used to place the 66kb-sized blocks, resulting in over 250k block scores.

#### 2.4.2 Principal Component Analysis (PCA)

The HapMap3 data set was used to calculate and plot the first two principal components from both variant based data and QGS based data. The QGS values were derived using HapMap3 itself as a reference set and the GENCODE gene annotation v3c release. Correlations between components were calculated. The full analysis scripts are available in the QGS repository.

#### 2.4.3 QGS distribution

The distribution of a metric can be important for statistical tests. The statistical distribution of QGS was demonstrated using two histograms based on (i) 1 variant and (ii) >1000 variants from UKBB-c.

#### 2.4.4 Effects of region size

A common problem for genetic aggregation methods is the influence of the size of the aggregated region on subsequent association (e.g. Liu et al., 2010). The influence of region size on QGS association was tested using a permutation approach using UKBB-c, NBS, and 1000G. The a priori distribution of region sizes (in this case, gene sizes) was compared to the distribution of region sizes in the top one-percent of associated genes from 1000 simulated binary phenotype permutations. QQ plots were made to visualise all three distributions and a Kolmogorov-Smirnov (K-S) test was used to quantify differences in distribution.

#### 2.4.5 Effects of LD structure

In addition to the effects of region size, genetic aggregation methods also have to account for inflation of results because of LD correlation structure. The effects of LD structure on QGS were tested in a similar way as the effects of region size above: 1000 associations based on permutation of a simulated binary phenotype resulted in LD strength distribution of top one-percent associated genes, which was compared to the a priori LD strength distribution. LD strength here was defined as the mean pairwise R^2^ for neighbouring SNPs within a gene.

#### 2.4.6 Effects of flanking regions

Important genetic information sometimes lies outside the immediate genetic regions of interest, for example in case of promotor and enhancer regions of genes (depending on the gene definition used). Moreover, a large portion of significant GWAS association results have been found in intergenic regions (Bartonicek *et al.*, 2017). Because of this, it is common to include flanking regions of varying size in region-based genetic analyses. The Pearson correlation between QGS without flanking regions and QGS including varying sizes of symmetrical flanking regions was examined in UKBB-c, NBS, and 1000G.

#### 2.4.7 Effects of LD pruning

LD pruning is the removal of highly correlated variants based on LD and is sometimes employed before genetic aggregation, for example in polygenic risk scoring applications. The effects of LD pruning on the Pearson product-moment correlation coefficient of QGS without pruning and with various pruning thresholds were tested in NBS and 1000G.

#### 2.4.8 Effects of imputation quality control

Imputation is a statistical technique used in genetics to assign values to unmeasured variants based on genotyped variants using LD structure (Li *et al.*, 2009). Every imputed variant carries a measure of quality control, generally expressed as an estimated correlation (R^2^) or info score between the imputed and the real, unknown genotype. Imputed variants with poor quality are often removed from subsequent analyses. The correlation between QGS values using different thresholds of imputation quality control were examined in NBS. Imputed data was used at four levels of imputation quality control to determine the effects of the quality control step: (a) the raw imputed variants (no quality control); (b) imputed variants with a threshold of R^2^> 0.3 (low quality control); (c) imputed variants with a threshold of R^2^> 0.6 (medium quality control); and (d) imputed variants with a threshold of R^2^> 0.9 (strict quality control).

#### 2.4.9 Estimated number of independent tests

In GWAS, each genetic variant can give rise to a hypothesis test. However, due to LD structure, the tests are not independent. The typical multiple-testing corrections performed in genetic association studies rely on an accurate estimate of the number of independent tests (Benjamini *et al.*, 2001). Therefore, the number of independent tests for QGS genes was estimated in NBS using the method by Li and Ji (Li and Ji, 2005).

### 2.5 Traditional association studies

Three association studies were performed using the following phenotypes and data: ADHD diagnosis in the iPSYCH data set (binary outcome), BMI in the NBS data set (quantitative trait outcome), lifetime cannabis use in UKBB-c (binary outcome), and sociability in UKBB-s (quantitative trait outcome). Sex, age (for iPSYCH and NBS only), and the first 20 principal components were included as covariates. To maximise coverage, all types of gene regions (including non-coding genes and pseudogenes) were included, as well as 66kb blocks as described in section 2.4.1. Results of QGS were compared to previously published GWAS results to study overlap and differences. The focus of the comparison lies with identifying which variant-based information is captured in both GWAS and QGS as well as where results differ.

### 2.6 Stability selection study

Because QGS reduces the number of variables in GWAS, it may have more statistical power to detect genetic associations in small samples by leveraging computationally intensive methods. To demonstrate, stability selection using randomized lasso (Meinshausen and Bühlmann, 2010) was performed using the same NBS BMI data set described above. The stability selection method uses a permutation-based LASSO approach which, compared to GWAS, is computationally expensive. It was selected as an example of more complex methods that may be performed after dimensionality reduction that works well for data sets with many variables (i.e. genes) with a limited sample size that are underpowered for traditional GWAS analysis.

In short, whereas the goal of GWAS is to find the strongest associations, the goal of stability selection with QGS is to find the most stable associations. Since individual genetic effect sizes for multifactorial diseases tend to be small, looking for the most stable effects instead of the largest effects can help identify novel candidates.

### 2.7 Gene set analyses

Genetic effects on multifactorial conditions are often found distributed across the genome (Cookson *et al.*, 2009). Whereas the previous section focused on single region, this section explores the combined effects of sets of genes. Here, we constructed gene sets based on previous association study findings. Alternatively, gene sets could also be defined based on other methods, e.g. pathway analyses or specific hypotheses.

#### 2.7.1 BMI gene set analysis

A BMI gene set QGS sum score was constructed using 48 BMI risk genes from (Locke *et al.*, 2015). The sum score was calculated by summing QGS values for all 48 risk genes together into a single variable. Two association tests were performed in NBS, one focussing on the continuous BMI outcome, and a second on the binary BMI>25 overweight outcome. Both association tests included age and gender as covariates, as well as the first 10 principal components.

#### 2.7.2 Lifetime cannabis use gene set analysis

A lifetime cannabis use QGS sum score was constructed using 38 risk genes from (Pasman *et al.*, 2018) and tested in UKBB-c. Like above, the sum score was calculated by summing up the QGS values from individual risk genes. Since these risk genes were found in a largely overlapping sample, they do depend on our UKBB-c data. The goal of this analysis is not to verify the risk genes, but instead to show that QGS can capture gene set based information. An association test was done using gender and the first 10 principal components as covariates.

## 3 Results & Discussion

First, the results from the PCA analysis will be presented. Second, the analytical properties of QGS will be shown. Third, GWAS results from iPSYCH, NBS, and UKBB will be presented. Fourth, the stability selection results from NBS will be set out. Last, results from the two gene set based analyses will be shown.

### 3.1 PCA

After quality control, the HapMap3 data included 458,572 variants. QGS reduced this number of variables by 95% to 21,792 genes (without guaranteeing the same coverage: around 40% of variants was wholly discarded because they lie outside of gene regions). The results from the QGS-based PCA analysis are shown in Figure 2. The variant-based results are close to identical to the QGS-based results, with high correlation r>0.99 between the components. The proportion of genetic variance explained differs somewhat: 9.1% for the variant-based first component compared to 6.6% for QGS. A larger version of Figure 2 and the related variant-based PCA plot can be found in supplemental Figure S1.

**Figure 2:**
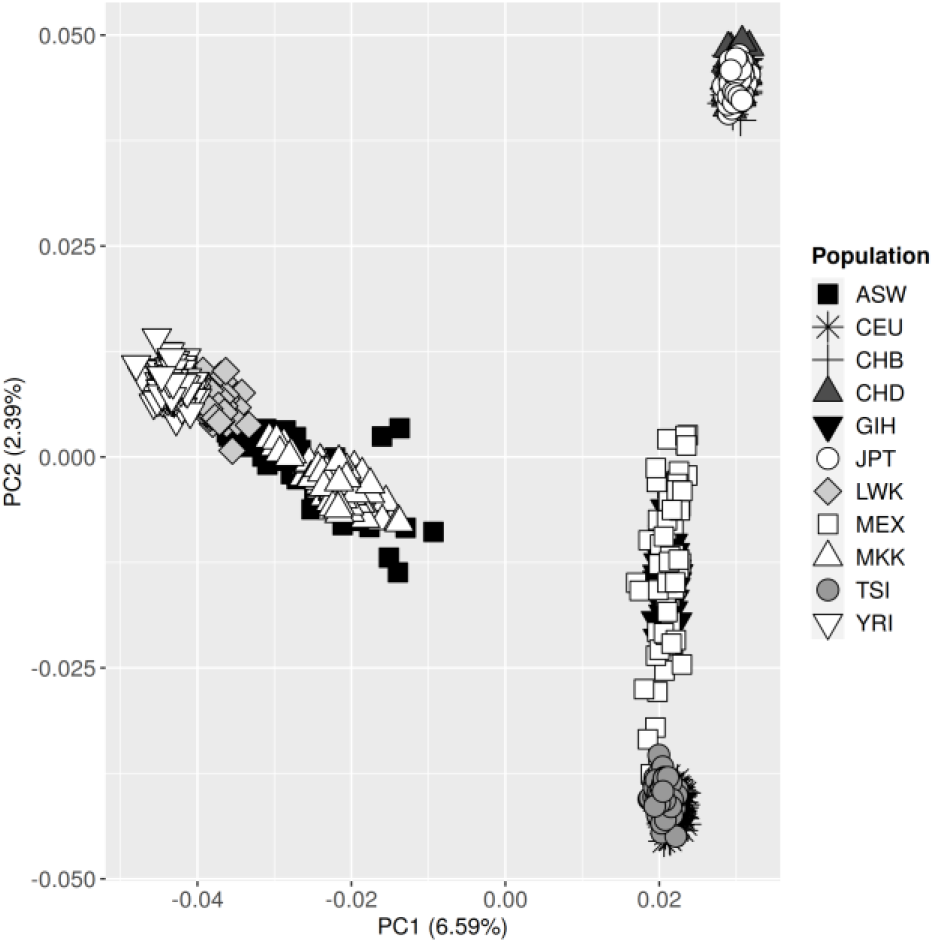
Principal component population stratification plot of 21,792 QGS values based on the genotypes of HapMap3 (N = 1,184). The axes show the first and second principal component (PC); the percentages represent the proportion of variance explained. ASW: African ancestry in Southwest USA; CEU: Utah residents with Northern and Western European ancestry; CHB: Han Chinese in Beijing, China; CHD: Chinese in Metropolitan Denver, Colorado; GIH: Gujarati Indians in Houston, Texas; JPT: Japanese in Tokyo, Japan; LWK: Luhya in Webuye, Kenya; MXL: Mexican ancestry in Los Angeles, California; MKK: Maasai in Kin-yawa, Kenya; TSI: Toscani in Italia; YRI: Yoruba in Ibadan, Nigeria.

### 3.2 QGS Properties

In this section, the analytical properties of QGS will be presented.

#### 3.2.1 Distribution of QGS

QGS scores based on a single called variant follow a binomial distribution with n=2, because there are three possible QGS values an individual can have based on a single variant: (a) one for individuals homozygous for the major allele, (b) one for heterozygous individuals, and (c) one for individuals homozygous for the minor allele. The distribution of these groups depends on the minor allele frequency (MAF) of the variant. In supplemental Figure S2A an example is given of the QGS distribution based on a single variant with low MAF. As the number of variants in a QGS value increases, this distribution approaches normality following the central limit theorem. This is illustrated in supplemental Figure S2B.

No clear guideline can be given for how many variants need to be included in a QGS value before it can be treated as approximately normal, because this depends on the MAF and LD of the included variants. In case of doubt or low amount of variation in a region, it is recommended to use statistical tests that do not depend on normality of the variable.

#### 3.2.2 Effects of region size and LD structure

The effects of gene size and LD structure on association results in three genetic data sets were examined. For both gene size and LD structure, no significant effects were observed, meaning that the associations based on QGS were not significantly affected by either of the two. For region size no effects were found in 1000G (p=0.14), NBS (p=0.99), or UKBB (p=0.96). Similarly, for LD structure no effects were found in 1000G (p=0.32), NBS (p=0.99), or UKBB (p=0.99). These findings are illustrated in supplemental Figures S3 and S4.

#### 3.2.3 Effects of flanking regions

Using symmetrical flanking regions of 5kb resulted in a correlation of 0.74-0.85 (with the QGS estimated without flanking regions) across the three data sets, whereas 25kb gives a correlation of 0.58-0.75, 50kb gives a correlation of 0.51-0.68, and 100kb 0.42-0.59. Since the mean size of a gene in our datasets is 66kb, a flanking region of 5kb on both sides increases the mean gene region size by about 15%, and 25kb increases the size by about 76%. The correlation is highest in the small NBS sample and lowest in the large UKBB-c sample. The full correlation matrices are shown in supplemental Figure S5.

#### 3.2.4 Effects of pruning

Since most closely neighbouring variants are highly correlated, the removal of all variants with a correlation threshold of R^2^>0.99 already resulted in a removal of 30-50% of variants from our data sets. The correlation between QGS without pruning and QGS with R^2^>0.99 pruning threshold is between r=0.91-0.93. At a pruning threshold of R^2^=0.9 through which more than half of the variants have been removed from the original data, the QGS correlation is r=0.83-0.88. The complete data is shown in supplemental Figure S6: (A) shows the effects of the pruning threshold on the QGS correlation and (B) shows the percentage of variants removed at each pruning threshold.

#### 3.2.5 Effects of imputation quality control

The correlation between QGS without quality control (R^2^ >= 0) and with (a) low quality control (R^2^ > 0.3) was r=0.99, (b) with medium quality control (R^2^<0.6) was r=0.97, and (c) with high quality control (R^2^>0.9) was r=0.89. The full correlation matrix is shown in supplemental Figure S7.

#### 3.2.6 Estimated number of independent tests

When testing a set of 32,221 known gene regions taken from NBS using a GWAS-like approach, the estimated number of independent tests was 24,263, or 76% of the total number of tests. This percentage was used subsequently in this paper to correct for the number of tests performed in QGS-based GWAS. The breakdown per chromosome can be found in supplemental Table S1.

### 3.3 Traditional association studies

#### 3.3.1 ADHD Case/control

A Manhattan plot for chromosome 1 containing both the variant-based GWAS and the QGS-based GWAS can be found in Figure 3. A genome wide Manhattan plot can be found in supplemental Figure S8 and a full overview of overlapping and differing regions can be found in supplemental Tables S2 and S3. The genome wide top hit in both analyses implicates the same region containing the SLC6A9 gene. The second-highest GWAS peak also contains the second-most significant QGS gene on this chromosome. This shows that information contained in QGS overlaps with variant-based information.

**Figure 3:**
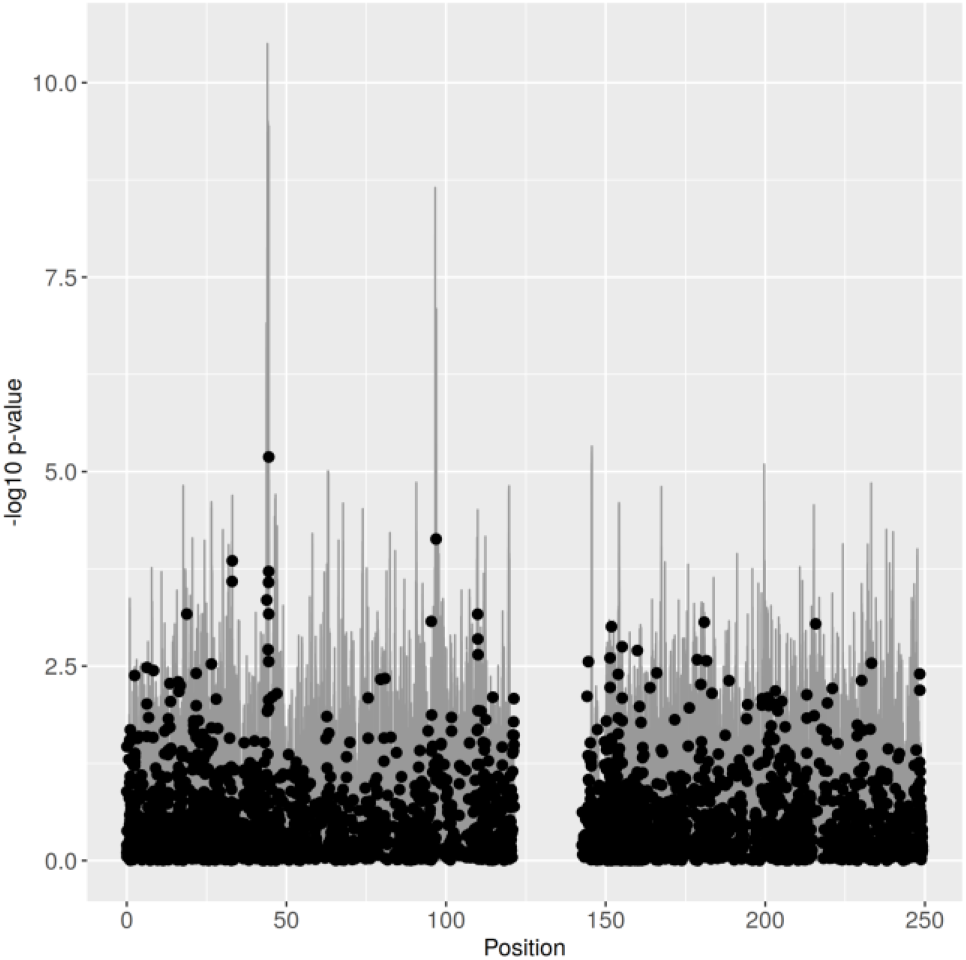
Manhattan plot of chromosome 1 for the variant-based and QGS_gene_-based association results of ADHD status. Negative log10 of the p-value is plotted on the y-axis and base pair position in million base pairs is plotted on the x-axis. The variant-based results are represented by grey bars. The QGS-based results are represented by black points.

Differences can also be found, for example the variant-based GWAS finds a significant variant on chromosome 16 contained in the LINC01572 gene that the QGS-based association does not identify. Vice versa, QGS-based association finds a suggestive result in a region on chromosome 7 containing the TRIM73 gene that the variant-based GWAS does not pick up. This suggests that individual variant-level associations can be lost in QGS. Overall, the highest variant-based peaks (top hit p=3.2e-11) have higher statistical significance than the QGS-based peaks (top hit p=6.5e-06).

#### 3.3.2 UKBB

##### 3.3.2.1 Lifetime cannabis use case/control

Similar to the ADHD results above, a Manhattan plot containing both the variant-based GWAS and the QGS-based GWAS for UKBB-c can be found in supplemental Figure S9. Whereas the variant-based GWAS finds significant results, the QGS-based GWAS does not. The top variant-based finding (rs28732378/3:85,403,892, p=3.4e-10) overlaps with top (but nonsignificant) block-based QGS results (rank 16 and 22). Apart from one other location on chromosome 4, little overlap is seen between the top variant-based results and QGS results. Conversely, the top QGS-based results do tend to have a low variant-based p-value. The top results from variant-based GWAS and QGS-based GWAS can be found in supplemental Tables S4 and S5.

##### 3.3.2.2 Sociability

Results in UKBB-s show a similar pattern as UKBB-c above. While the top variant-based GWAS hit is among the top 100 QGS-based results, the overlap between top variant-based results and QGS-based results is low (supplemental Table S6). Again and vice versa, top QGS-based results do tend to have a low variant-based p-value (supplemental Table S7). The full results can be seen in supplemental Figure S10.

#### 3.3.3 BMI GWAS in NBS

A GWAS analysis was performed for BMI in the smaller NBS sample. The variant-based GWAS identified one genome-wide significant variant (15:98876579/rs184852001, p=1.9e-08), but this SNP is not found in the largest BMI meta-GWAS to date (closest variant downstream rs1372847 with p=1.7e-01 and closest upstream rs4965808 with p=8.7e-01) (Yengo *et al.*, 2018). Neither the gene-based nor the block-based QGS finds significant results in this data set. QGS reduced the number of variables tested by 99.8% from 20,011,335 variants to 35,550 genes. Overlapping findings include regions on chromosome 5 (TMEM173) and 19 (ENSG00000267448). A Manhattan plot containing both the variant-based GWAS and the QGS-based results can be found in supplemental Figure S11 and a full overview can be found in supplemental Tables S8 and S9.

### 3.4 BMI stability selection

In contrast to the GWAS approach described in section 3.3.3, the stability selection with QGS yields stable results for BMI in the smaller (n=4,454) NBS data set, as shown in Figure 4. The three stable genes are VSX1, GON7, and EPGN. Each of these has (or contains variants that have) previously been associated with obesity (Locke *et al.*, 2015; Fox *et al.*, 2007; Winkler *et al.*, 2015; Akiyama *et al.*, 2017; Rankinen *et al.*, 2006). Moreover, the first two genes have been replicated in the largest BMI meta GWAS to date (VSX1 with rs6138482, p= 5.8e-13 and GON7 with rs7148516, p=1.3e-17) (Yengo *et al.*, 2018).

**Figure 4:**
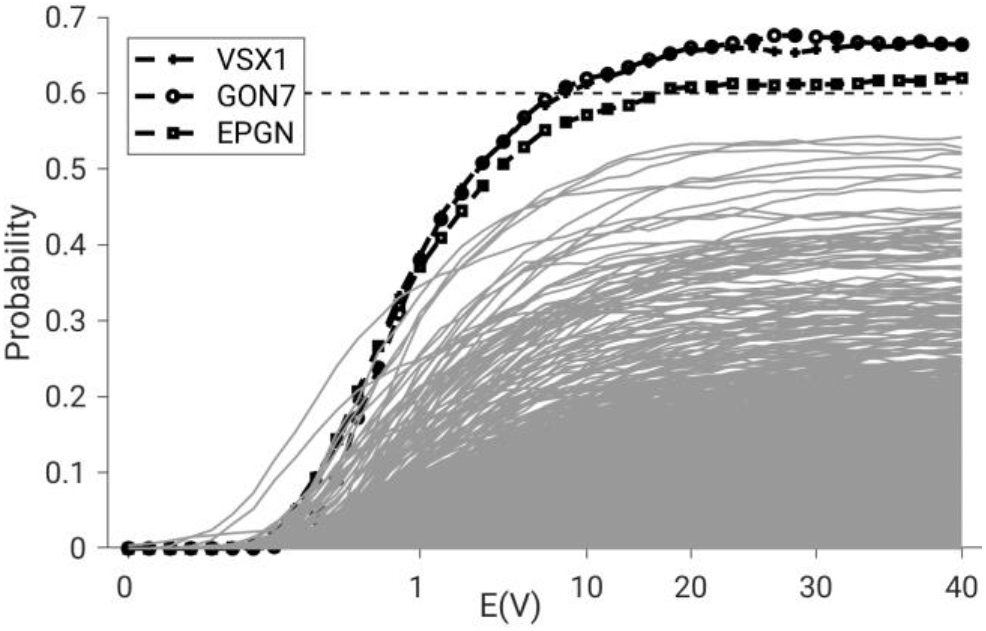
Stability selection results for BMI using NBS; N = 4,454, N_variant_ = 8,476,119, N_QGS-gene_ = 18,942 (protein coding genes only). The E(V) value on the x axis represents the upper limit for the expected number of false positives (see Meinshausen and Bühlmann, 2010) and is a function of the strength of the LASSO penalty: the more relaxed the penalty, the higher E(V). The selection probability on the y axis represents the probability that a gene is included in the LASSO-selected set for the current E(V)/LASSO penalty. The y=0.6 dotted line represents our cut-off point for stability.

### 3.5 Gene set analyses

#### 3.5.1 BMI gene set

The QGS sum of the BMI gene set consisting of 48 obesity risk genes is associated with BMI (p=.026, b(se)=32.7(14.7), 95%CI 3.8/61.6) as well as with an obesity construct consisting of BMI>25 (p=.021, b(se)=18.1(7.9), 95%CI=2.7/33.6) in NBS. The QGS sum explained 0.46-0.48% of BMI variance in the NBS sample. Associations including every gene in the gene set univariately as well as all together (in a multiple regression) did not provide any multiple-testing-corrected significant results, the top hit being the gene VKORC1 (p=.014). These results are shown in supplemental Table S10.

#### 3.5.2 Lifetime cannabis use gene set

A lifetime cannabis use QGS sum score was constructed using 38 risk genes. This sum score was associated with lifetime cannabis use (p=0.024, b(se)=-0.022(0.01), 95%CI −0.042/−0.003). The QGS sum score explained 0.006% of cannabis use variance in the UKBB sample. The regression results can be found in supplemental Table S11.

## 4 Conclusion

This work introduced QGS, a method to create meaningful quantitative variables for arbitrary genetic regions. Using complex phenotypes in differently sized real samples, QGS was shown to reduce the number of variables with 98-99% yet conserve relevant information at low, medium, and high level of granularity using respectively PCA, gene sets, and individual genes. In combination with stability selection, it was shown to be successful in correctly identifying relevant genes where traditional GWAS was not.

At low level of granularity, PCA analysis shows that QGS retains the same rough population-level demographic information that can be found in variant-level data.

At a medium level of granularity, gene set analyses show that a QGS value for a selected set of genes can explain a small but significant portion of phenotypical variance for BMI and cannabis use. This shows that QGS can highly compress genetic information - in this case, to a single number for 40-50 genes - and still capture phenotypic variance. The gene set score based on QGS is agnostic to the direction of effect, so while it is likely that not all genes in the gene sets have the same direction of effect, this does not prevent us from finding significant effects.

At a high level of granularity, stability selection results show that the combination of QGS with more complex variable selection methods can yield results in cases where traditional GWAS-like screening struggle. In addition, the three stable and verified genes show that - despite a 99.8% reduction in variables - QGS retains relevant genetic information for complex phenotypes at the gene level.

Limitations of QGS include information loss at the single variant level because of the significant dimensionality reduction, shown in the moderate overlap between variant-based and QGS-based GWAS. In addition, QGS appears sensitive to the effects of data pruning, where the removal of highly correlated variants leads to a proportionally larger reduction in QGS correlation. Because pruning (if performed) is done data set wide, this sensitivity does not affect association results. In contrast, QGS is insensitive to factors that have previously been shown to affect association, such as region size and LD structure, and it is stable when including flanking regions.

Because QGS is phenotype-agnostic, it only needs to be calculated once. After that, it can be used in any downstream analysis for any available phenotype. Applications include the use of genes, gene sets, or genetic regions in regressions, in common statistics software such as SPSS, or more complex methods like stability selection, random forests, or causal discovery. By reducing genetic sequencing data to a single construct of arbitrary resolution, QGS enables the combination of genetic information with other information sources into complex models. QGS is available at https://github.com/machine2learn/QGS.

## Acknowledgements

This work was supported by the European Union’s Seventh Framework Programme for research, technological development and demonstration under grant agreement no 602805 - AGGRESSOTYPE; and the research programme Computing Time National Computing Facilities Processing Round pilots 2018 with project number 17666, which is (partly) financed by the Dutch Research Council (NWO). This work was partially carried out on the Dutch national e-infrastructure with the support of SURF Cooperative. This paper reflects only the author’s views and the European Union is not liable for any use that may be made of the information contained therein.

