## Supplemental figures for "Quantitative Genetic Scoring, or how to put a number on an arbitrary genetic region"

**Figure S1**: Principal component population stratification plot of (A) 21,792 QGS values on the left and (B) 458,572 variants (SNPs) on the right based on the genotypes of HapMap3 (N = 1,184). The axes show the first and second principal component (PC); the percent-ages represent the proportion of variance explained. ASW: African ancestry in Southwest USA; CEU: Utah residents with Northern and Western European ancestry; CHB: Han Chinese in Beijing, Chin a; CHD: Chinese in Metropolitan Denver, Colorado; GIH: Gujarati Indians in Houston, Texas; JPT: Japanese in Tokyo, Japan; LWK: Luhya in Webuye, Kenya; MXL: M exican ancestry in Los Angeles, California; MKK: Maasai in Kinyawa, Kenya; TSI: Toscani in Ita-lia; YRI: Yoruba in Ibadan, Nigeria.

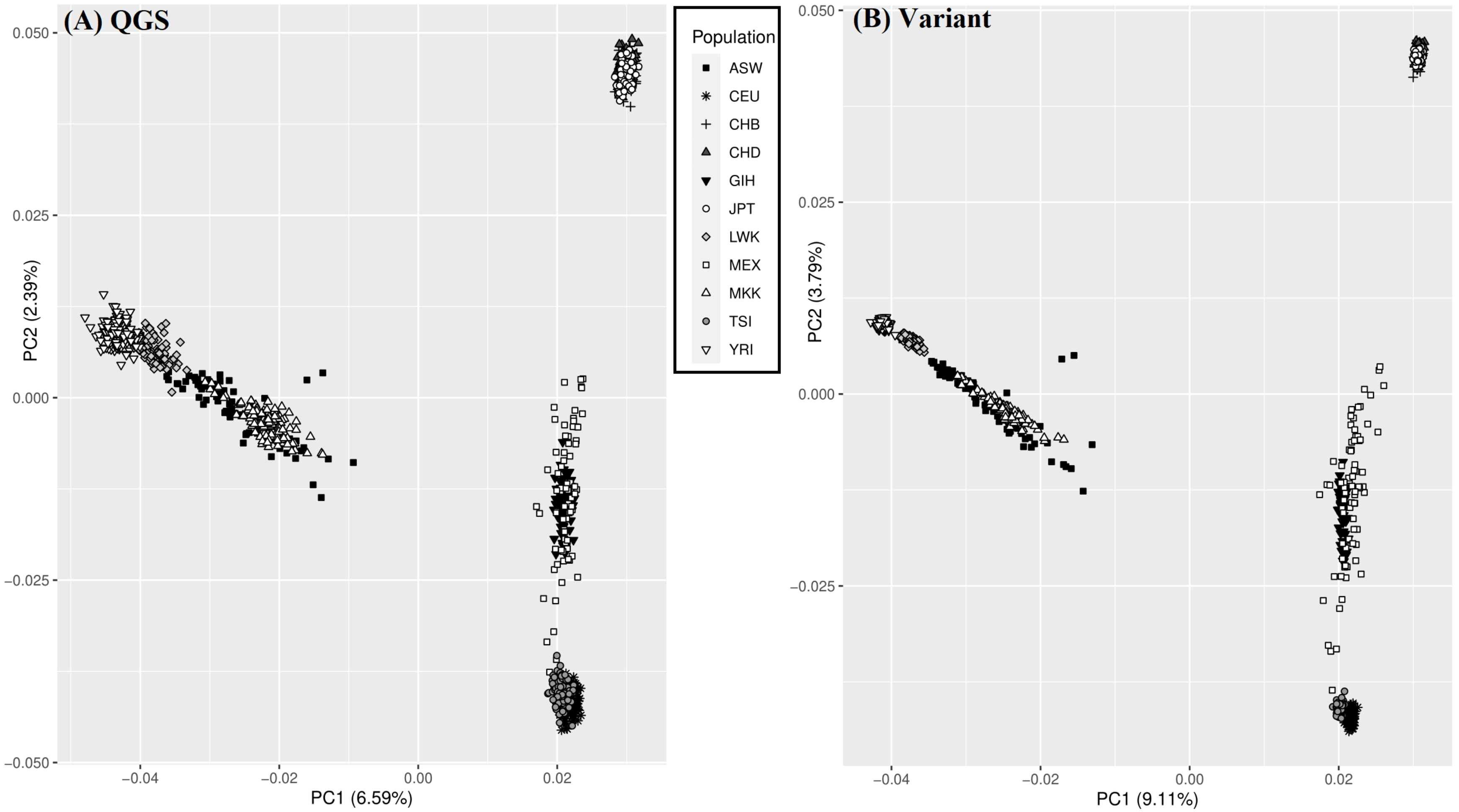

**Figure S2**: Distribution of quantitative genetic score for genes ($QGS_{gene}$) for a coding gene containing 1 variant and a coding gene containing 1,069 variants using UKBB-c (N = 153,501) as sample and reference. These two genes were selected to demonstrate the effects of the number of variants on a $QGS_{gene}$ distribution. The single variant in the HIST1H4G gene is rs41266821 (MAF=0.12 in UKBB-c).

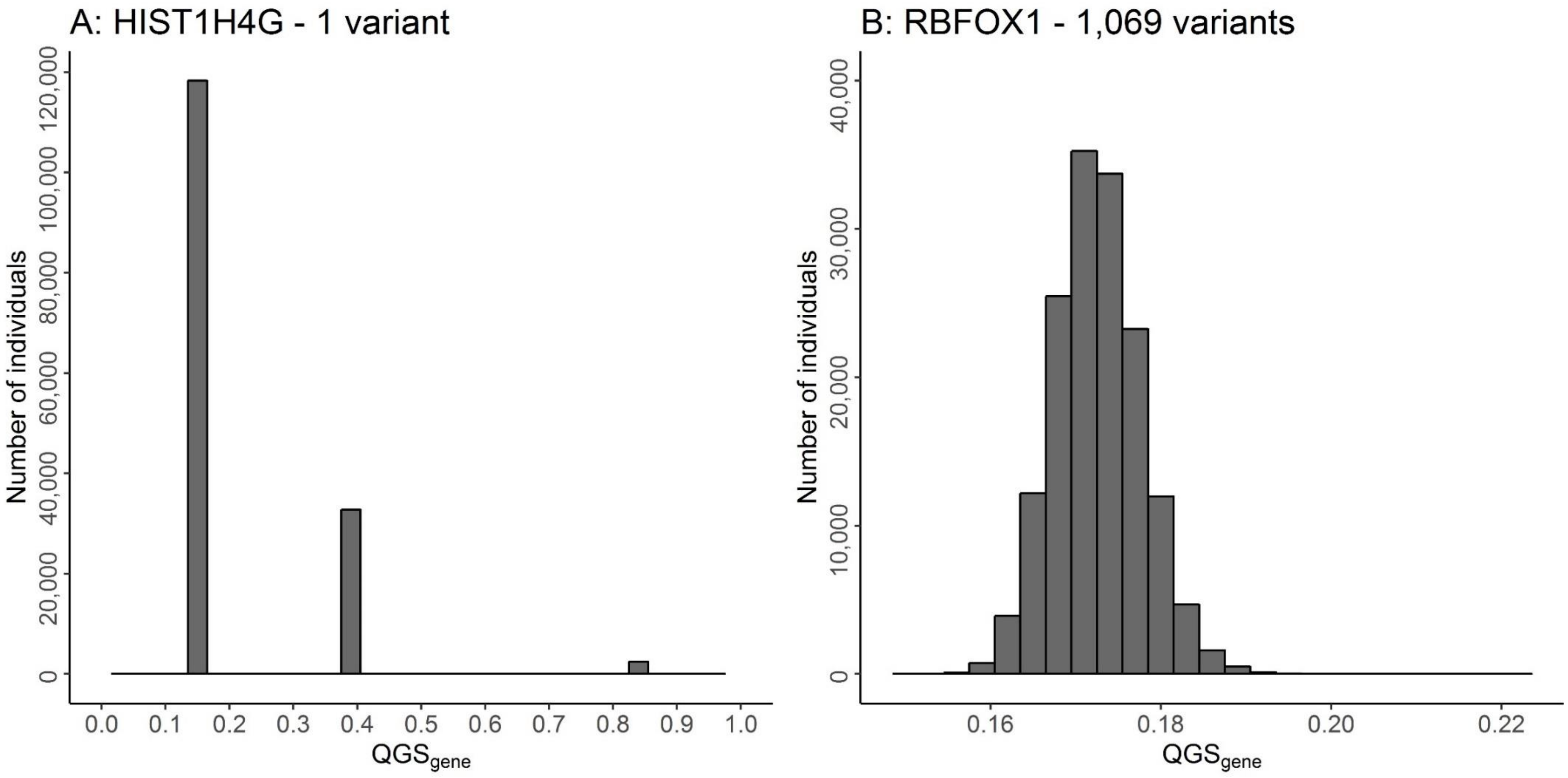

**Figure S3**: Effects of region size on $QGS_{gene}$-based association results using genotypes from 1000genomes database (A: 1000g; N = 2,504; $N_{QGS-gene}$ = 49,669), Nijmegen Biomedical Study (B: NBS; N = 4,454; $N_{QGS-gene}$ = 45,372) and UK-Biobank database (C: UKBB; N = 153,501; $N_{QGS-gene}$ = 29,751). Quantile-Quantile (Q-Q) plots of the expected region size distribution (x-axis) against the observed region size distribution of top 1% associated genes (y-axis) using 1,000 simulated associations per gene based on a random binary phenotype. The dotted line denotes the pattern under the null hypothesis. No significant differences in distribution were observed (K-S: Kolmogorov-Smirnov test).

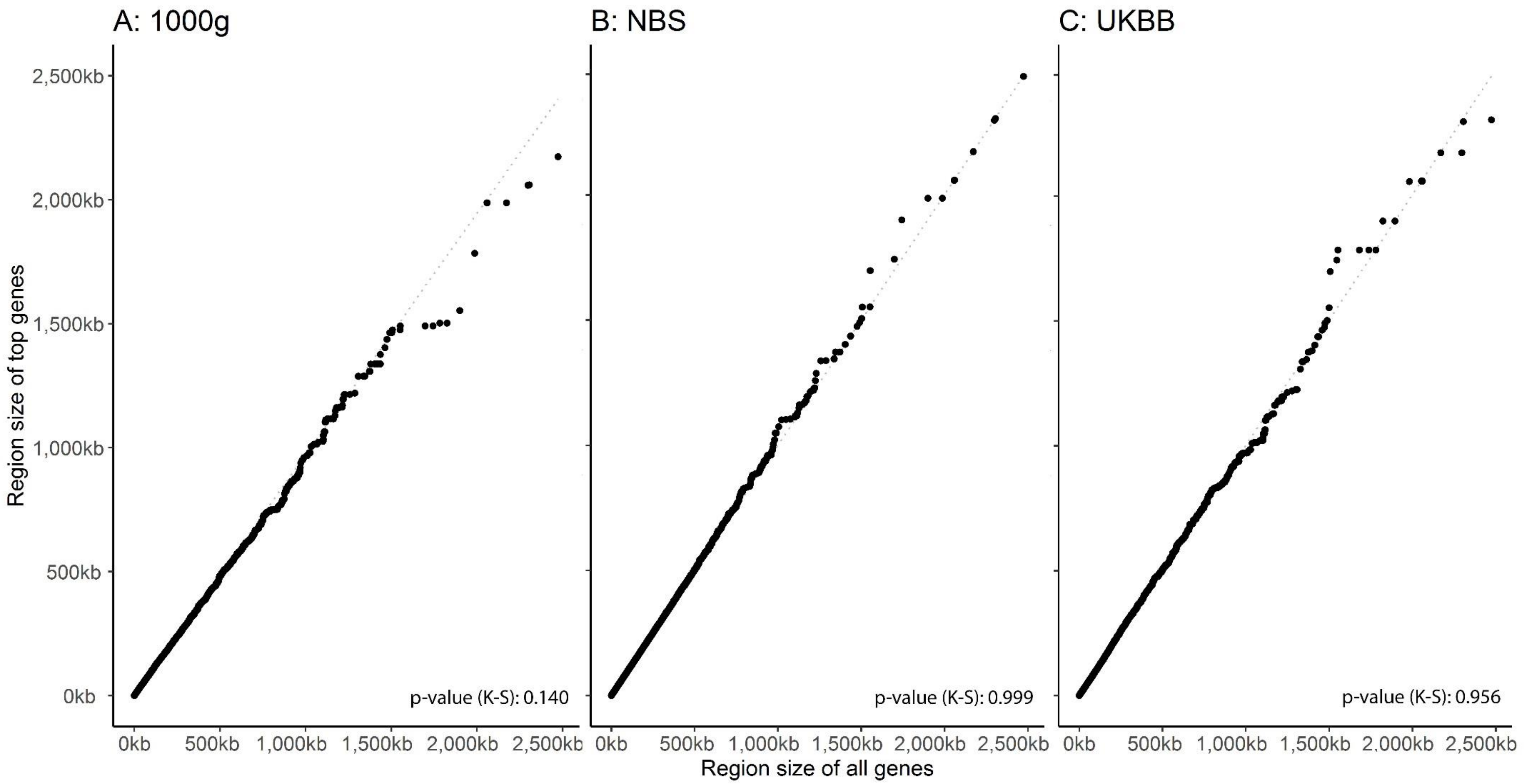

**Figure S4**: Effects of LD structure on QGSgene-based association results using genotypes from 1000genomes database (1000g; N = 2,504; $N_{QGS\text{-}gene}$ = 49,669), Nijmegen Biomedical Study (NBS; N = 4,454; $N_{QGS\text{-}gene}$ = 45,372) and UK-Biobank database (UKBB; N = 153,501; $N_{QGS\text{-}gene}$ = 29,751). Quantile-Quantile (Q-Q) plots of the expected LD structure distribution (x-axis) against the observed LD structure distribution of top 1% associated genes (y-axis) using 1,000 simulated associations per gene based on a random binary phenotype. The dotted line denotes the pattern under the null hypothesis. No significant differences in distribution were observed (K-S: Kolmogorov-Smirnov test).

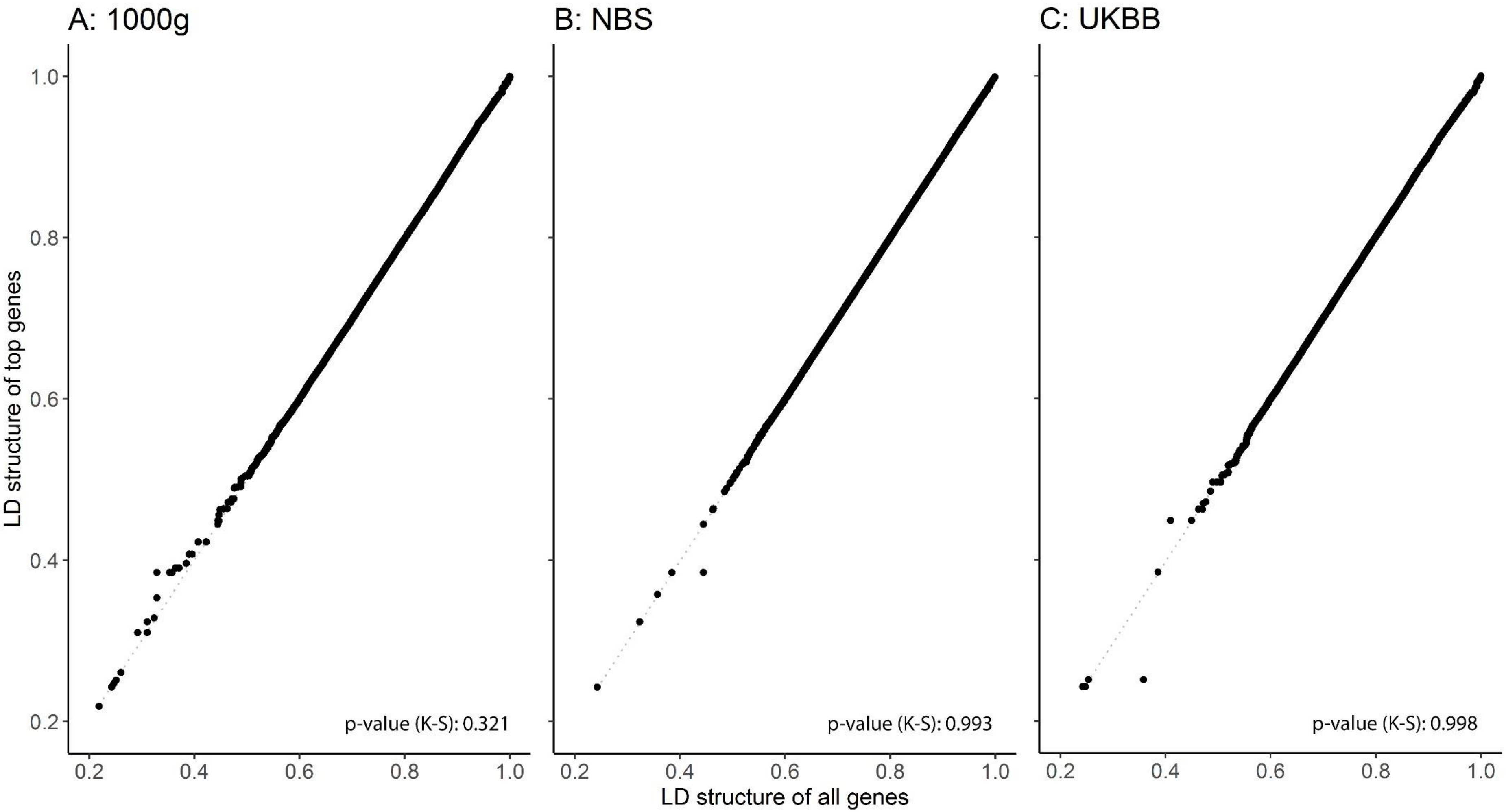

**Figure S5**: Effects of flanking on $QGS_{gene}$. Correlation matrix of $QGS_{gene}$ allowing for 0kb flanking region to $QGS_{gene}$ allowing for the current flanking region using genotypes from 1000genomes database (A: 1000g; N = 2,504; $N_{QGS\text{-}gene}$ = 49,669), Nijmegen Biomedical Study (B: NBS; N = 4,454; $N_{QGS\text{-}gene}$ = 45,372) and UK-Biobank database (C: UKBB-c; N = 35,000 (randomly selected subsample); $N_{QGS\text{-}gene}$ = 29,751). Five flanking settings were used: a flanking region of 0kb, a flanking region of 5kb, a flanking region of 25kb, a flanking region of 50kb and a flanking region of 100kb.

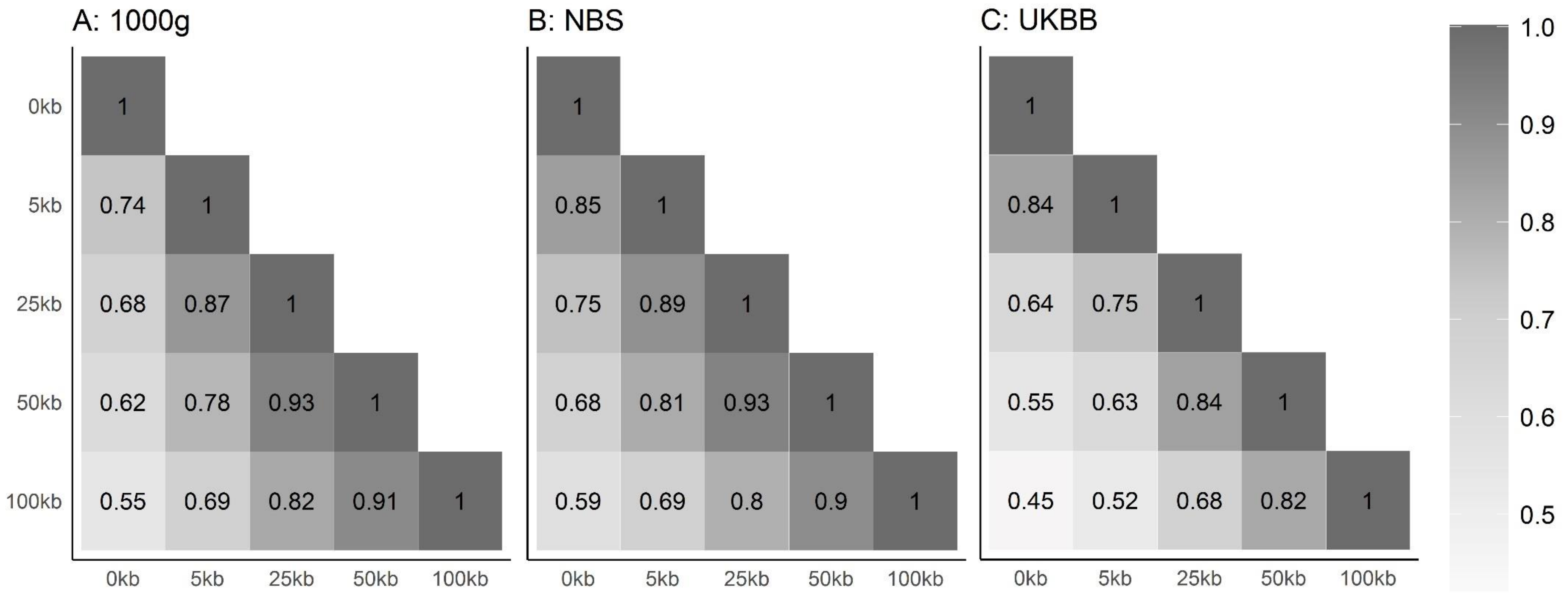

**Figure S6**: Effects of pruning on quantitative gene score of genes ($QGS_{gene}$). Correlation matrix of $QGS_{gene}$ of no pruning with the current pruning threshold using genotypes from 1000genomes database (1000g; N = 2,504), Nijmegen Biomedical Study (NBS; n = 4,454) and UK-Biobank database (UKBB; N = 153,501). Twenty-four pruning thresholds were used from no pruning threshold to a pruning threshold of 0.05 (0.05:0.95 with steps of 0.05 and 0.96:1.00 with steps of 0.01). The QGSgene is calculated based on the number of coding genes found in the dataset with a pruning threshold of 0.05 leading to NQGS -gene = 13,694 for 1000g, NQGS-gene = 7,681 for NBS and NQGS-gene = 6,429 for UKBB.

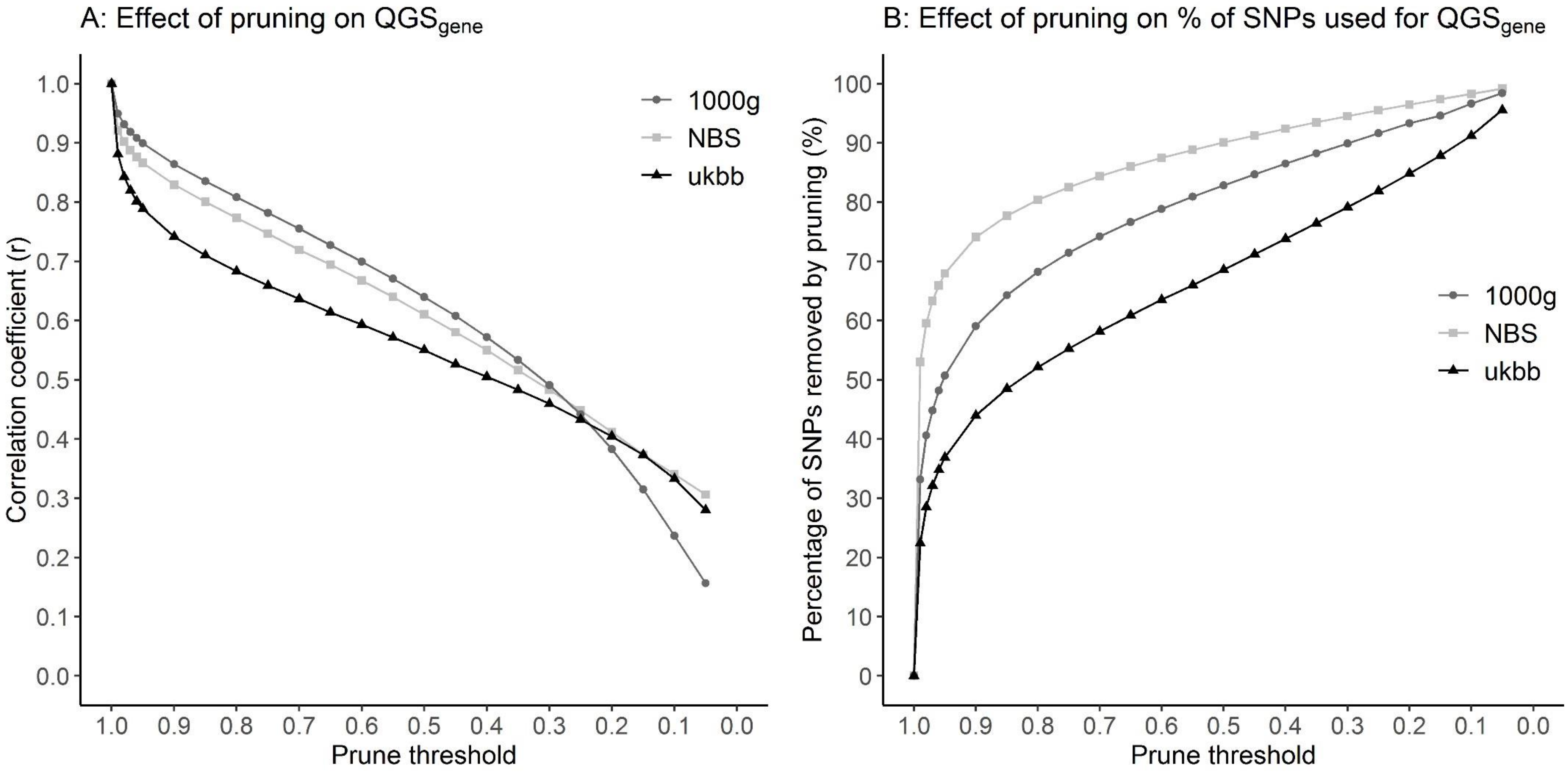

**Figure S7**: Effect of imputation quality control on $QGS_{gene}$. Correlation matrix of $QGS_{gene}$ of $R^2 \geq 0$ with the current imputation quality using genotypes from Nijmegen Biomedical Study (NBS; N = 4,454; $N_{QGS\text{-}gene}$ = 45,372). Four imputation qualities were used: no imputation (R2 ≥ 0), an imputation quality of 0.3 (R2 > 0.3), an imputation quality of 0.6 (R2 > 0.6) and an imputation quality of 0.9 (R2 > 0.9). The QGSgene is calculated based on: 27,303,679 SNPs for the R2 ≥ 0 row; 9,472,350 SNPs for the R2 > 0.3 row; 7,132,038 SNPs for the R2 > 0.6 row and 4,387,293 SNPs for the R2 > 0.9 row.

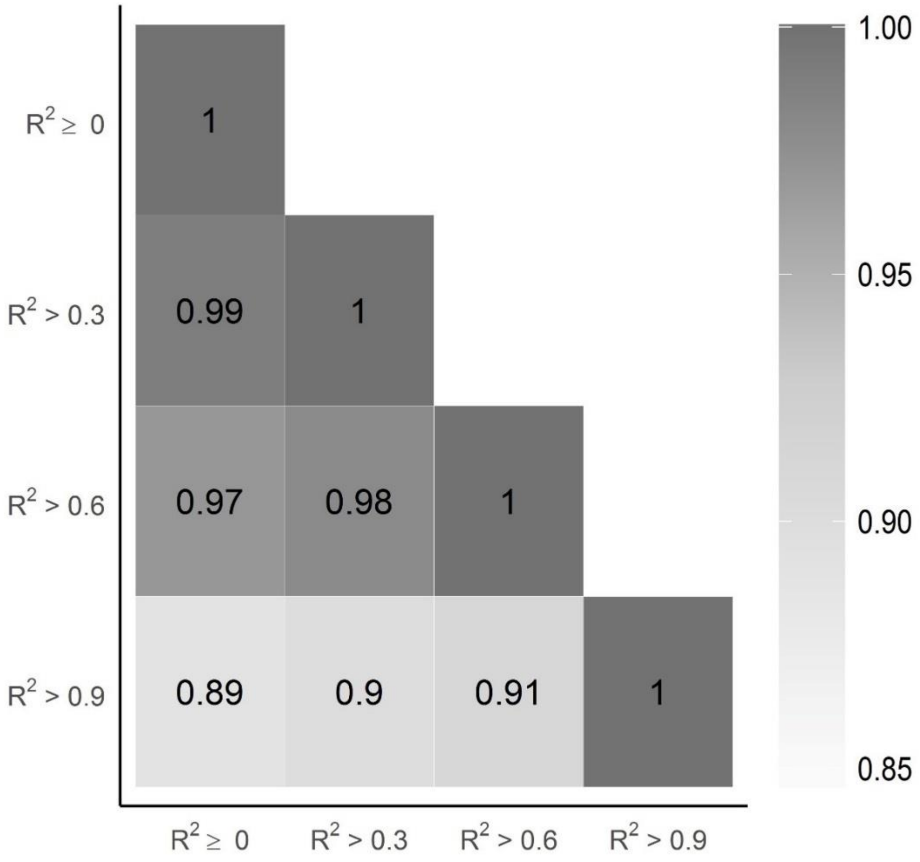

**Table S1: Calculation of independent tests when using quantitative genetic score of genes ($QGS_{gene}$) in a univariate model using genotypes from Nijmegen Biomedical Study (NBS; n = 4,452).**

| CHR | Number of genes | Number of independent tests | Percentage |
|---|---|---|---|
| 1 | 3,128 | 2,266 | 72 |
| 2 | 2,290 | 1,762 | 77 |
| 3 | 1,740 | 1,332 | 77 |
| 4 | 1,580 | 1,290 | 82 |
| 5 | 1,703 | 1,356 | 80 |
| 6 | 1,904 | 1,442 | 76 |
| 7 | 1,702 | 1,275 | 75 |
| 8 | 1,332 | 1,067 | 80 |
| 9 | 1,287 | 1,002 | 78 |
| 10 | 1,327 | 1,042 | 79 |
| 11 | 2,063 | 1,420 | 69 |
| 12 | 1,731 | 1,294 | 75 |
| 13 | 786 | 666 | 85 |
| 14 | 1,127 | 873 | 77 |
| 15 | 976 | 707 | 72 |
| 16 | 1,271 | 870 | 68 |
| 17 | 1,707 | 1,143 | 67 |
| 18 | 659 | 566 | 86 |
| 19 | 1,917 | 1,349 | 70 |
| 20 | 886 | 689 | 78 |
| 21 | 412 | 339 | 82 |
| 22 | 693 | 515 | 74 |
| Total | 32,221 | 24,263 | 76 |

The number of genes is the total number of genes per chromosome with a QGS value. In a univariate model, this is the number of individual tests to perform. Since these tests will not be totally independent because of the LD structure between genes, the number of independent tests is calculated whilst controlling for dependency. The percentage shows the ratio of number of genes and number of independent tests.

**Figure S8**: Manhattan plot for the variant-based and $QGS_{gene}$-based association results of ADHD. − log10 (p-value) is plotted on the y-axis and chromosomal location is plotted on the x-axis. The variant-based results are represented by grey bars. The $QGS_{gene}$-based results are represented by the green. The dash-dotted line represents the conventional genome-wide significance threshold of p-value < 5e-08. The dotted line represents a suggestive significance level of p-value < 1e-06. The dashed line represents a suggestive significance level of p-value < 1e04.

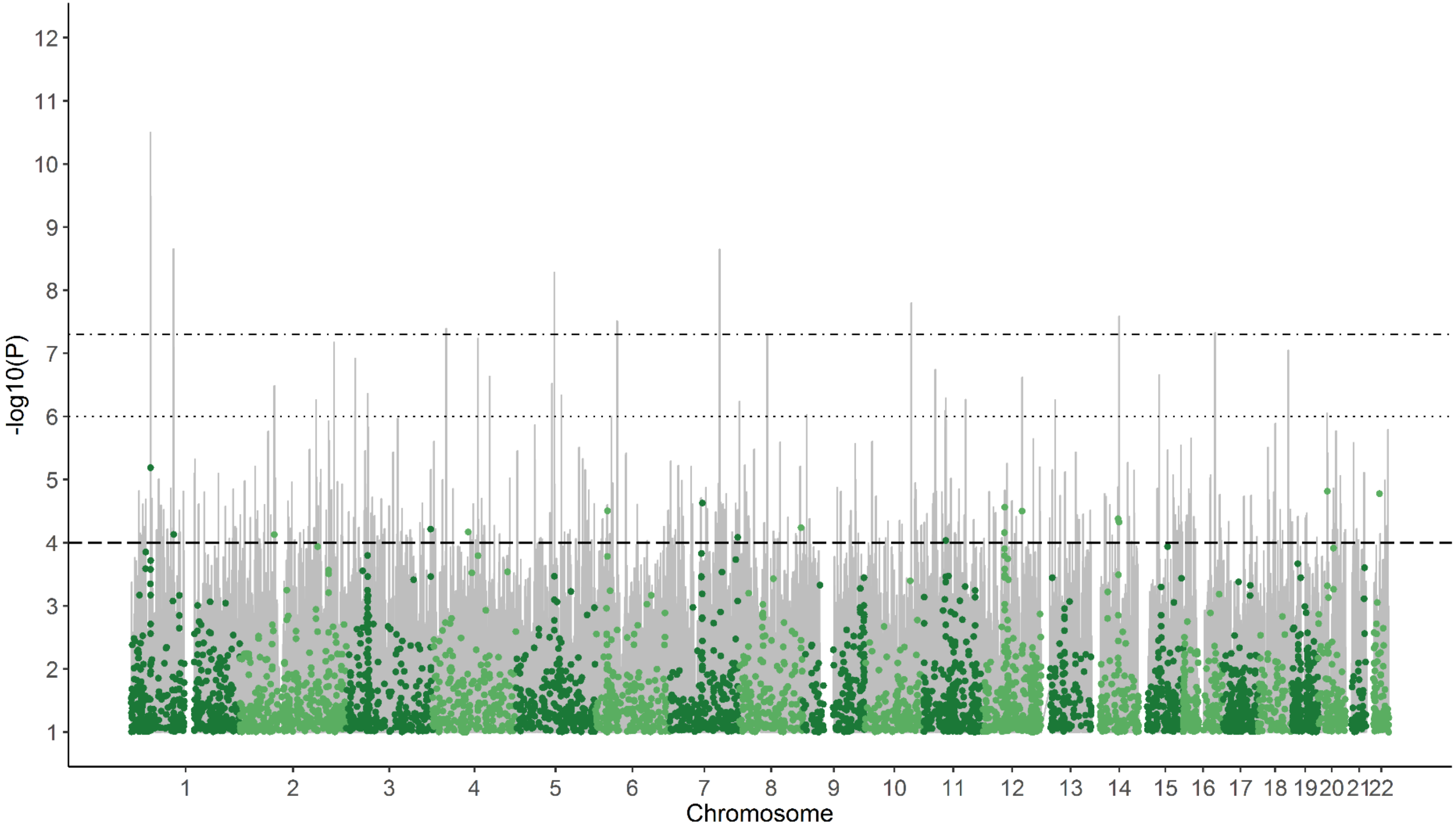

**Figure S9**: Manhattan plot for the variant-based, $QGS_{gene}$-based and $QGS_{block}$ association results of cannabis use. Results are based on data from UKBB-c; N = 131,864, $N_{variant}$= 14,021,094, $N_{QGS\text{-}gene}$ = 29,751 and $N_{QGS\text{-}block}$.=264,111. The − log10 (p-value) is plotted on the y-axis and chromosomal location is plotted on the x-axis. The variant-based results are represented by grey bars. The $QGS_{gene}$-based results are represented by the green dots and the $QGS_{block}$-based results are represented by the purple dots. The dash-dotted line represents the conventional genome-wide significance threshold of p-value < 5e-08. The dotted line represents a suggestive significance level of p-value < 1e-06. The dashed line represents a suggestive significance level of p-value < 1e04.

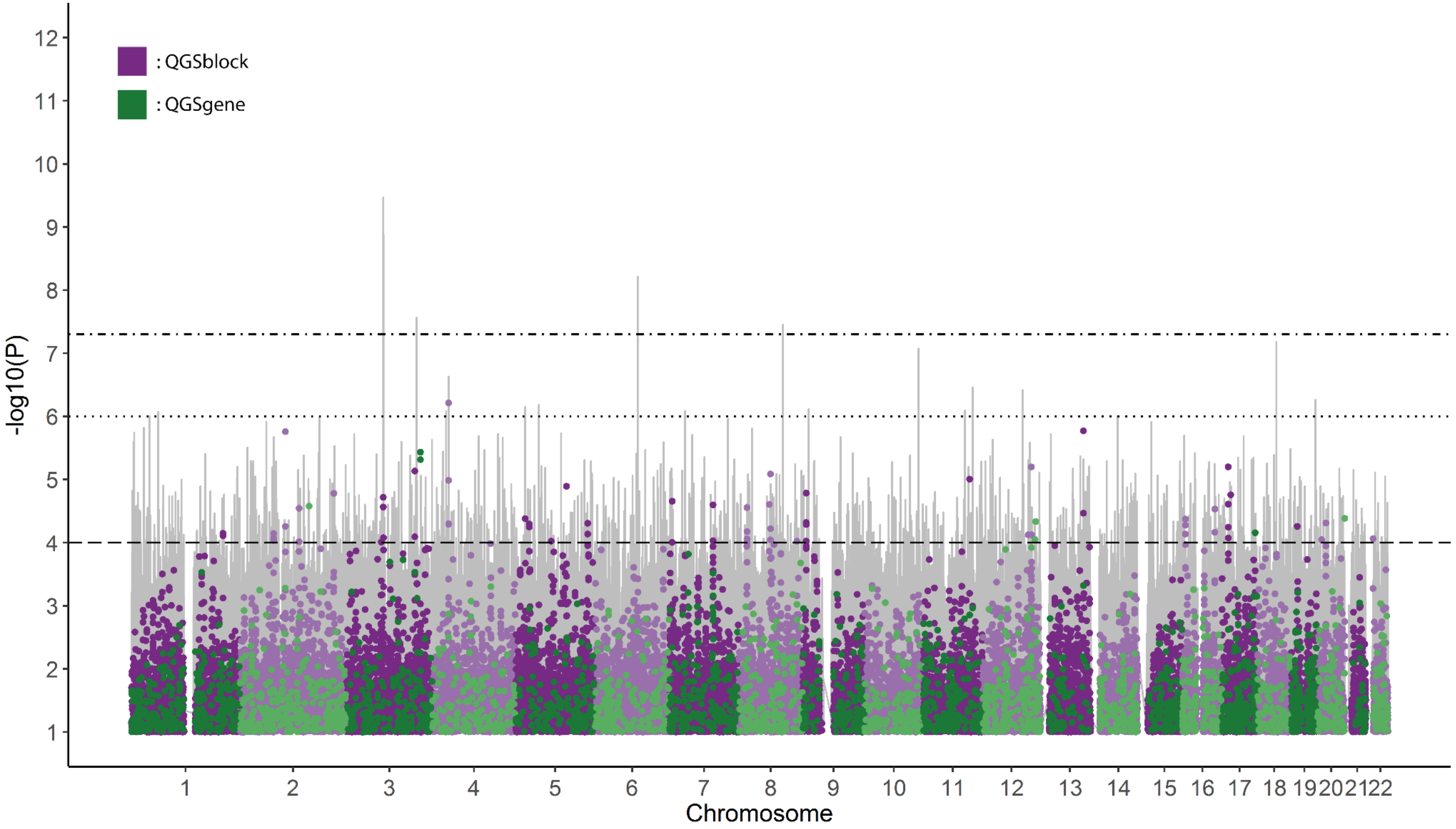

**Figure S10**: Manhattan plot for the variant-based, $QGS_{gene}$-based and $QGS_{block}$-based association results of sociability. Results are based on data from UKBB-s; N = 342,461; $N_{variant}$= 10,903,884; $N_{QGS\text{-}block}$ = 264,187; $N_{QGS\text{-}gene}$ = 30,016. The -log10 (p-value) is plotted on the y-axis and chromosomal location is plotted on the x-axis. The variant-based results are represented by grey bars. The $QGS_{gene}$-based results are represented by the green dots and the $QGS_{block}$-based results are represented by the purple dots. The dash-dotted line represents the conventional genome-wide significance threshold of p-value < 5e-08. The dotted line represents a suggestive significance level of p-value < 1e-06. The dashed line represents a suggestive significance level of p-value < 1e04.

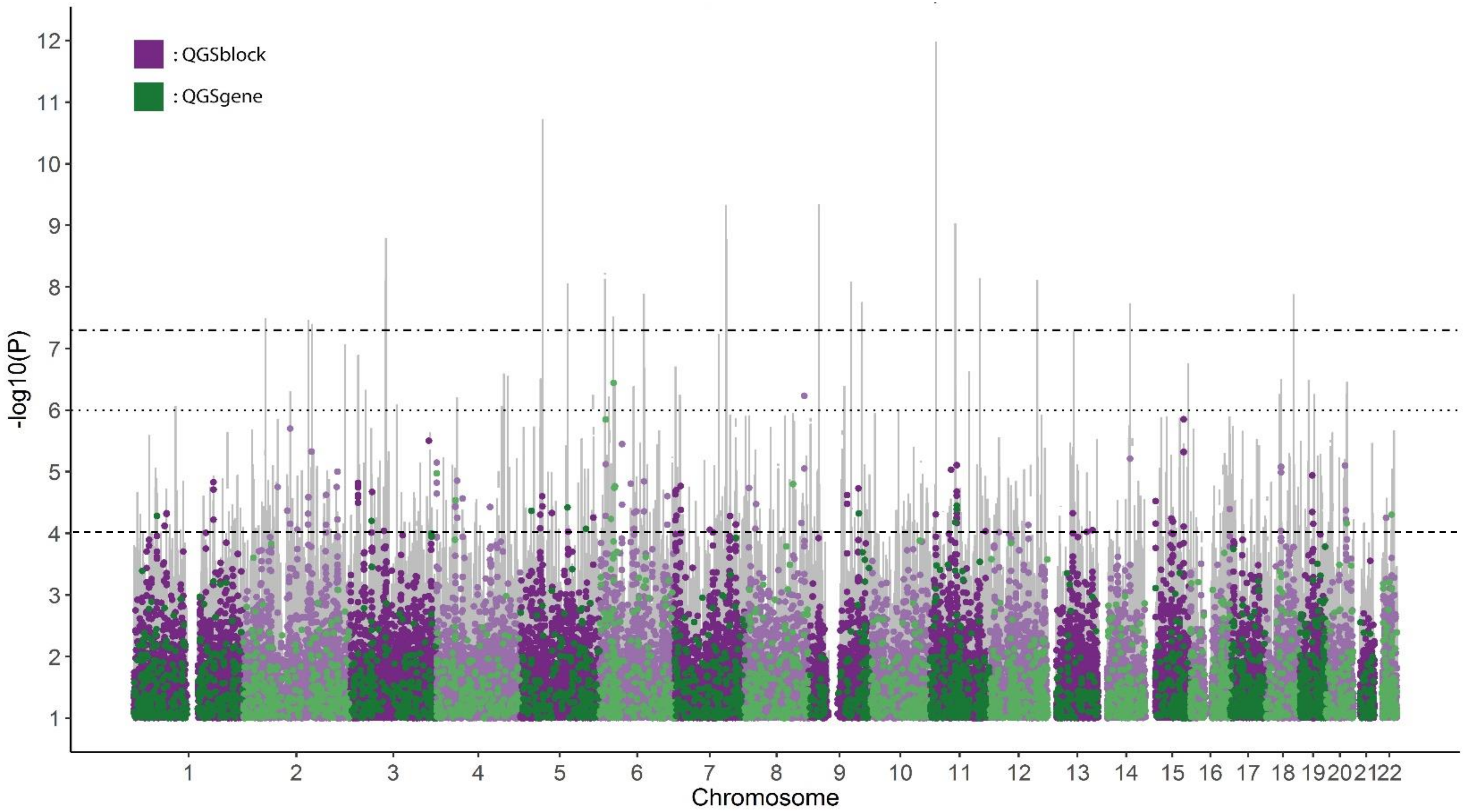

**Figure S11**: Manhattan plot for the variant-based, $QGS_{gene}$-based and $QGS_{block}$-based association results of BMI. Results are based on data from NBS; N = 4,454, $N_{variant}$ = 8,476,119, $N_{QGS-gene}$ = 45,372 and $N_{QGS-block}$ = 267,622. The variant-based results are represented by grey bars. The $QGS_{gene}$-based results are represented by the green dots and the $QGS_{block}$-based results are represented by the purple dots. The dash-dotted line represents the conventional genome-wide significance threshold of p-value < 5e-08. The dotted line represents a suggestive significance level of p-value < 1e-06. The dashed line represents a suggestive significance level of p-value

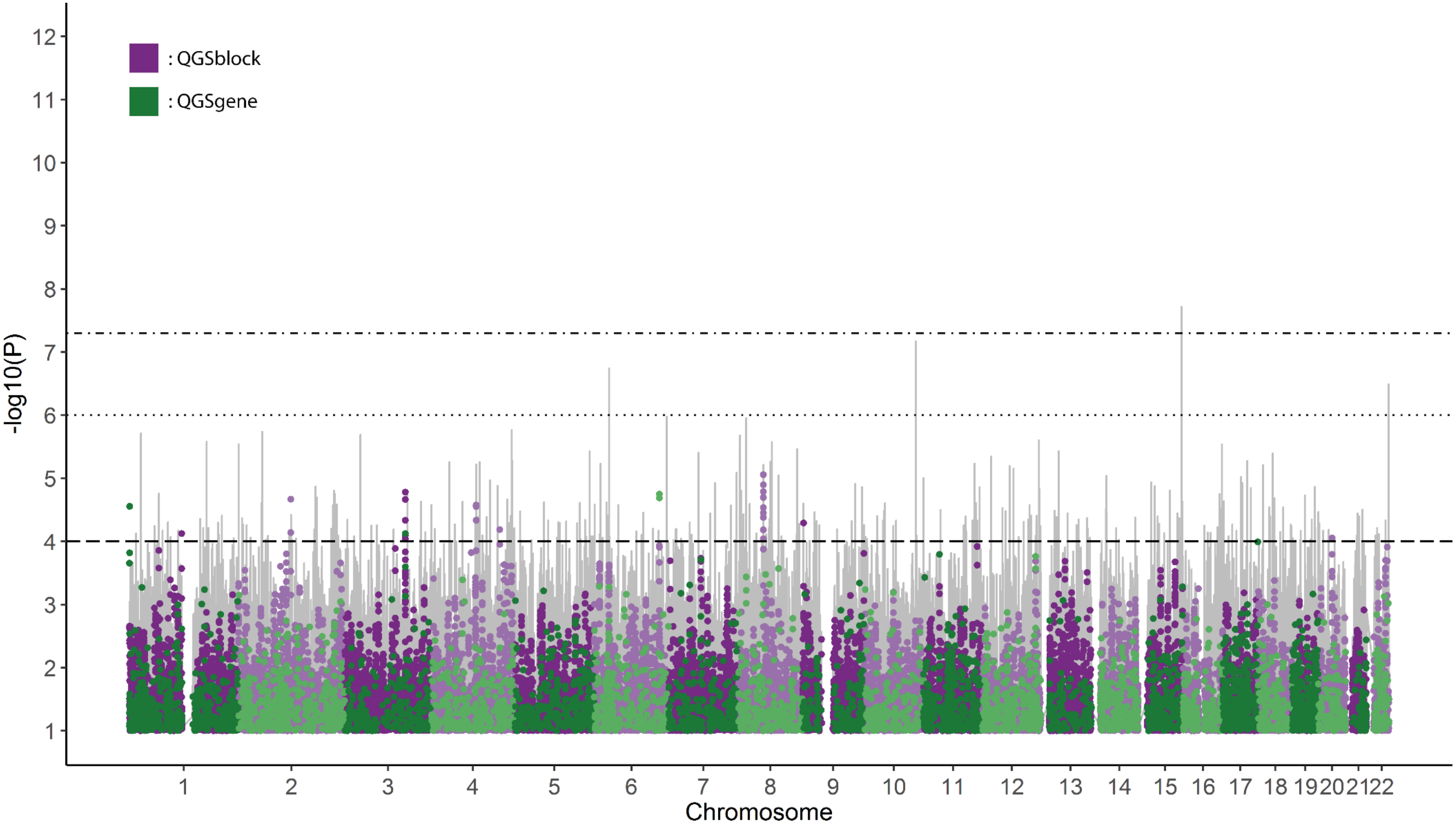


< 1e04.

**Tables S2-S9** are included as Excel tables in a separate document.

**Table S10**: Results for the association of the sum quantitative genetic score of 47 obesity risk genes ($\sum QGS_{gene}$) and Body Mass Index (BMI) as linear variable or as logistic variable (BMI > 25 and BMI < 25). Results are based on data from NBS; N = 4,454 and $N_{variant}$= 8,518,806.

| | B (SE) | CI | P-value | $R^2$ % |
|---|---|---|---|---|
| $\sum QGS_{gene}$ linear | 32.721 (14.714) | 3.847 – 61.596 | 2.639e-02* | 0.460 |
| $\sum QGS_{gene}$ logistic | 18.106 (7.873) | 2.656 – 33.56 | 2.147e-02* | 0.480 |

The regression model is corrected for age, gender and principal components 1-10; B: regression coefficient; S.E.: Standard Error; CI: 95% Confidence Interval; $R^2$%: percentage of variance explained. The statistical significance is marked with asterisks for *p-value<0.05.

**Table S11** Results for the association of cannabis use and the sum quantitative genetic score of 38 genes linked to lifetime cannabis ($\sum QGS_{gene}$). Results are based on data from UKBB; N = 131,864 and $N_{variant}$= 14,021,094.

| | B (S.E.) | 95% CI | P-value | $R^2$ % |
|---|---|---|---|---|
| $\Sigma$ $QGS_{gene}$ | -0.022 (0.010) | -0.042 - -0.003 | 2.400e-02* | 0.006 |

The regression model is corrected for gender and principal components 1-10. B: regression coefficient; S.E.: Standard Error; CI: 95% Confidence Interval; $R^2$%: percentage of variance explained.

### QGS estimation example

As an example, consider a genetic region **s** defined by 2 SNPs: rs1 and rs2. For a single sample individual, the dosage vector is given by s = {0.2, 1.2}, where 0.2 is the estimated (imputed) dosage for rs1 and 1.2 is the estimated imputed dosage for rs2. A reference population consisting of 3 individuals is defined as matrix

$$R = \begin{bmatrix} 0.5 & 0.8 \\ 1.1 & 0.1 \\ 1.4 & 0.3 \end{bmatrix}$$

where the top left 0.5 represents the estimated imputed dosage for rs1 for reference individual 1, etc. Following Equation 1 for the QGS, the absolute difference between the sample individual and each reference individual is computed for every SNP:

$$QGS = \sum_{r=1}^{3} \sum_{i=1}^{2} |R_i^r - s_i|$$

$$= |0.5 - 0.2| + |0.8 - 1.2| + |1.1 - 0.2| + |0.1 - 1.2| + |1.4 - 0.2| + |0.3 - 1.2|$$

$$= 0.3 + 0.4 + 0.9 + 1.1 + 1.2 + 0.9 = 4.8$$

Then the summed result is scaled to a value between 0-1:

$$QGS = \frac{\sum_{r=1}^{N_{refs}} \sum_{i=1}^{N_{snps}} |R_i^r - s_i|}{2N_{refs}N_{snps}} = \frac{4.8}{2 \cdot 3 \cdot 2} = \frac{4.8}{12} = 0.4$$

This final result of 0.4 represents the QGS for the sample individual calculated with the provided reference. Note that the QGS does not depend on phenotype and as such does not need to be recalculated for different applications, in contrast with existing genetic aggregation methods such as weighted PRS.

When the reference panel, the included variants, and the QGS value are all known, a limited amount of genetic information from $\vec{s}$ can be inferred. From the above example, one could e.g. deduce that the estimated imputed dosage for rs2 for the sample individual must be ≥1 by solving the below for rs2.

$$|0.5 - rs1| + |0.8 - rs2| + |1.1 - rs1| + |0.1 - rs2| + |1.4 - rs1| + |0.3 - rs2| = 4.8$$

This deduction becomes increasingly less informative with the inclusion of more SNPs in the genetic region, because the number of unknown variables increases.

### Genetic Data collection and processing

*UKBB*

The data collection and processing in UKBB has been extensively described (Bycroft *et al.*, 2018; Welsh *et al.*, 2017). In short, retrieval and genotyping of blood samples was performed using a custom Affymetrix pipeline. Poor quality samples and markers were excluded, and imputation was performed with IMPUTE4 using the Haplotype Reference Consortium, UK10K, as well as 1000G as reference panels.

For UKBB-c data was made available by Pasman et al. and is described in (Pasman *et al.*, 2018). For UKBB-s data was made available by Bralten et al. and is described in (Bralten et al., 2019; preprint).

*iPSYCH*

For iPSYCH, data has been made available by Demontis et al. and is described in (Demontis *et al.*, 2019). In short, blood samples were sequenced using Illumina's Beadarrays and imputed using IMPUTE2 with the 1000G reference panel. Quality control steps included removing related individuals, genetic outliers, and filtering for MAF < 0.01.

*NBS*

For NBS, blood samples were collected and genotyping was performed using Illumina HumanOmniExpress-12 and -24 BeadChip genotyping platforms and imputation was performed using the 1000G reference panel. Quality control steps included removing related individuals, genetic outliers, filtering for MAF < 0.01, HWE < 1e-4, sex check, sample call rate, and a SNP yield > 95%. A more detailed description can be found in the original publication (Galesloot *et al.*, 2017).

*1000G*

1000G has been described in (Consortium *et al.*, 2010; 1000 Genomes Project Consortium *et al.*, 2015). In short, 1000G was genotyped fully; most variants were called using single read mapping, but multiple procedures were used with special provisions made for low coverage regions. Phasing was done using (custom) IMPUTE2 as well as BEAGLE algorithms and imputation-based techniques were used to produce the final genotype call and phasing information. For our analyses, no further processing was done.

*HapMap3*

Genotyping of blood samples was performed on Illumina (earlier) and Perlegen (later) platforms. Quality control steps differed per platform, but include HWE filtering, a maximum genotype missingness of five percent, and a Mendel errors threshold. In addition, SNPs without a valid rsID or unique genomic location were removed. More details can be found in the original publications (Altshuler *et al.*, 2010; Frazer *et al.*, 2007).

1000 Genomes Project Consortium,T. 1000 G.P. *et al.* (2015) A global reference for human genetic variation. *Nature*, **526**, 68–74.

Altshuler,D.M. *et al.* (2010) Integrating common and rare genetic variation in diverse human populations. *Nature*, **467**, 52–58.

Bralten,J. *et al.* (2019) Genetic underpinnings of sociability in the UK Biobank. *bioRxiv*, 781195.

Bycroft,C. *et al.* (2018) The UK Biobank resource with deep phenotyping and genomic data. *Nature*, **562**, 203–209.

Consortium,T. 1000 G.P. *et al.* (2010) A map of human genome variation from population scale sequencing. *Nature*, **467**, 1061.

Demontis,D. *et al.* (2019) Discovery of the first genome-wide significant risk loci for attention deficit/hyperactivity disorder. *Nat. Genet.*, **51**, 63–75.

Frazer,K.A. *et al.* (2007) A second generation human haplotype map of over 3.1 million SNPs. *Nature*, **449**, 851–861.

Galesloot,T.E. *et al.* (2017) Cohort Profile: The Nijmegen Biomedical Study (NBS). *Int. J. Epidemiol.*, **46**, 1099-1100j.

Pasman,J.A. *et al.* (2018) GWAS of lifetime cannabis use reveals new risk loci, genetic overlap with psychiatric traits, and a causal effect of schizophrenia liability. *Nat. Neurosci.*, **21**, 1161–1170.

Welsh,S. *et al.* (2017) Comparison of DNA quantification methodology used in the DNA extraction protocol for the UK Biobank cohort. *BMC Genomics*, **18**, 26.
